# Heat-tolerant network hub species make freshwater microbial communities resilient to climate warming

**DOI:** 10.1101/2025.06.06.658219

**Authors:** Wensi Zhang, Bharat Manna, Naresh Singhal

## Abstract

Aquatic ecosystems are facing severe threats from climate change, with rising temperatures as a major driving force posing a significant challenge to their ecological balance. Microorganisms constitute the majority of biomass in the water ecosystems and mediate the flux of carbon, nitrogen, sulfate and other essential nutrients. Climate warming profoundly influences microbial communities by shaping their distribution and ecological roles in the ecosystem, and in turn microorganisms play a significant role in climate feedback. The Yellow River, China’s second-longest river, is crucial for agricultural irrigation and provides drinking water for millions, yet little is known about microbial community adaptations under future warming scenarios in the system. In this study, we analyzed a publicly available metagenomic dataset from Yellow River water samples that were previously subjected to temperature treatments of 23, 26, 29, 32, and 35 degrees Celsius. We employed metagenomic analysis to identify how gradually increasing temperatures affected microbial community profiles. The study identified 140 species tolerant to high temperatures, showing a significant increase in abundance with elevated temperatures. The elevated temperature stress impacted the network properties of microbial communities substantially. Certain temperature-tolerant species were identified as hubs in the network across five temperatures, with their number increasing with temperature. Deltaproteobacteria bacterium was present at all five temperatures, while *Parcubacteria bacterium, Parvularcula sp*., *Phenylobacterium sp*., *Phycisphaeraceae bacterium*, and *Sphingobium xenophagum* were only present at high temperatures of 32 °C and 35 °C. The percentage of taxa nodes connected to tolerant hubs in the overall network rose from 57.48% to 95.94%, indicating the growing importance of these tolerant hubs. The positive connections among these tolerant hubs also increased, with the number of positive edges rising from 1166 to 2811, indicating a potential collaborative relationship among these taxa in response to temperature stress. These findings suggest that tolerant hubs may initially respond to temperature stress and subsequently transfer this function to other species through cooperative interactions. Understanding these microbial dynamics is crucial for developing strategies to maintain freshwater ecosystem health amid climate change.

## 1. Introduction

Climate change is widely regarded as the most severe challenge for both humanity and ecosystems. It has changed terrestrial, marine, and freshwater ecosystems all over the world, and impacts are widespread with more far-reaching consequences than anticipated (Pecl et al., 2017). A report from the Intergovernmental Panel on Climate Change (IPCC) in 2022 indicates that the situation has become even worse, with 3.3 billion of the world’s population exceptionally vulnerable to climate change (Jones, 2022). The worldwide observation from 2230 localities in 137 countries suggested that the global biodiversity decline was associated with increased temperature and changes in seasonality (Savo et al., 2016). A study in tropical waters showed that an increase in water temperature of 1-2 °C may result in severe mortality of phylum Cnidaria in reef-building corals (Reusch, 2014). Microorganisms constitute the majority of biomass on the earth, and climate warming profoundly influences microbial communities by shaping their distribution and ecological roles in the ecosystem (Singh et al., 2018). It was reported that warming could influence the prokaryotic community by largely remaining oligotrophic microbes in marine water (Yeh and Fuhrman, 2022). In turn microorganisms play a significant role in climate feedback as they impact the global ecosystem substantially by mediating the flux of carbon, nitrogen, sulfate and other essential nutrients (Hawkes et al., 2015; Hutchins et al., 2017). As a response to the increase in temperature, the distribution of phytoplankton communities has been observed to change globally, resulting in subsequent alterations in primary production (Poloczanska et al., 2013). In addition, some toxin-producing phytoplankton that are preferred in warming oceans may multiply rapidly and cause catastrophically detrimental algal blooms (Hutchins et al., 2019). Some microbes could even adapt to an extremely high-temperature environment. *Epsilonproteobacteria* constituted the most abundant community of bacteria in deep-sea hydrothermal habitats (Nakagawa et al. 2005). Microbes have developed a variety of functional metabolic strategies to adjust to high temperatures. Certain microorganisms were reported to reduce membrane fluidity through alterations in the lipid composition of cell membranes and increase the expression of heat shock proteins to respond to higher temperatures (Zheng et al., 2019). Another study on bacterioplankton from the South China Sea showed that their functional genes related to dissimilatory nitrate reduction, organic compounds degradation, DNA repair systems and biosynthesis of unsaturated biofilm were up regulated significantly (Ren et al., 2022). Compared to the ancestor, high-temperature-adapted Escherichia coli had higher N:P ratio but did not vary significantly in C:N and C:P ratios. The model inferred that adapted cells would produce a more massive amount of nitrogen-rich protein and phosphorus-rich ribosome in higher temperatures compared to their ancestor (Linzner et al., 2018).

Managing climate change challenges requires an in-depth understanding of intricate interactions between the climate system and ecosystems. Metagenomics especially can recover full genomes from environmental microbes and further profile the taxonomic composition and functional potential of microbial communities. This technology can enlarge our genomic knowledge of microorganisms in response to climate warming. Due to the critical functions of the microbes in climate warming responses, understanding how they will collectively adapt to changes is a vital goal (Hawkes et al., 2015). One study reported that warming significantly increased microbial network size, connectivity, and the number of central species. For example, it could trigger dynamic changes among different microbes by selecting certain fast-growing bacteria rather than slow-growing species (Mengting et al., 2021). Another study showed that the rise in temperature enhanced the resilience of the soil bacterial community network and boosted the capacity of bacteria to break down organic matter, thereby accelerating the soil carbon cycle (Yu et al., 2021). However, a 5-year experimental warming in a coastal soil ecosystem indicated a decrease in the complexity of the prokaryotic network, along with a simplification of the fungal-prokaryotic network (Zhou et al., 2021). Despite progress in elucidating the complexity of biological systems, many gaps remain in how warming stress affects the interaction among microbial communities.

Understanding how climate change will alter microbial communities and in turn how they will feed back to impact the pace of climate change requires substantial investigations in the ecosystem. Aquatic systems could support a variety of human activities, including water supply, fishing, recreation, and transportation, with rivers being especially significant. The Yellow River is the second-longest river in China and the sixth-longest river in the world (M. Yu et al., 2019). It holds a dominant position in agricultural irrigation with coverage of nearly 12% of the national area and serving almost 50 cities in northern China (Yu et al., 2022). The Yellow River provides drinking water for more than 4 million residents of Lanzhou city, making its water quality vital to the health of those living along its bank (Zhao et al., 2018). In this study, water from the Yellow River was heated to temperatures of 23 °C, 26 °C, 29 °C, 32 °C, and 35 °C to investigate the impact of elevated temperature on the microbial functions and interactions. The objectives of the study are to (1) Identify microbial species in the Yellow River that exhibit tolerance to elevated temperatures; (2) Examine the ecological roles of temperature-tolerant species in shaping microbial community structure, interactions, and functional processes under warming scenarios. We combine the experimental and metagenomics approach to understand how multilevel microbial processes change with elevated temperature, which may help us to anticipate possible consequences of global climate warming.

## 2. Methods

### 2.1. Sample data collection and metagenomics analysis

In this work, we analyzed shotgun metagenomics data from Yellow River water samples that were previously collected from the Lanzhou section (36°N, 103°E) and subjected to controlled temperature treatments of 23°C, 26°C, 29°C, 32°C, and 35°C (Yu et al., 2023). In the original experimental study, DNA was extracted and sequenced from each sample after one month of culture retention. The raw metagenomic data were publicly available under accession number PRJEB54031 (http://www.ebi.ac.uk/ena/data/view/PRJEB54031), and we analyzed these data using the SqueezeMeta v1.5.2 pipeline (Tamames & Puente-Sánchez, 2019). Raw sequences were first trimmed and filtered using Trimmomatic v0.39 (Bolger et al., 2014) for adapter removal, trimming, and quality control with specific quality thresholds. Subsequently, Megahit v1.2.9 (Li et al., 2015) was used for the assembly process to assemble raw reads into contigs. Prodigal v2.6.3 (Hyatt et al., 2010) was then applied for predicting open reading frames (ORFs) and retrieving corresponding amino acid sequences, while Barrnap v0.9 (Seemann, 2014) was used for RNA prediction. For taxonomic assignment, Diamond v2.0.14 (Buchfink et al., 2014) was used for homology searching against the NCBI GenBank nr database, with identity thresholds of 85, 60, 55, 50, 46, 42, and 40% for species, genus, family, order, class, phylum, and kingdom ranks, respectively. After taxonomic and functional assignment of contigs, all genes and contigs were mapped back to their respective samples. The metagenomics data was analyzed using the SQMtools v1.6.3 R package (Uritskiy & DiRuggiero, 2019). CSS (Cumulative Sum Scaling) normalized values were used to address compositional bias in microbiome data, indicating the abundance of taxa and genes (Paulson et al., 2013).

### 2.2. Network construction

We constructed networks for five temperature levels by using the NetCoMi R package (Peschel et al., 2021). Raw read count data were filtered to remove low-abundance taxa and subjected to zero replacement followed by centered log-ratio transformation to address compositional constraints (Martín-Fernández et al., 2003). SparCC associations between pairs of taxa were computed, considering coefficients higher than 0.80 or lower than -0.80, with a significance level of p < 0.05 (Tsilimigras & Fodor, 2016). The sparsified association matrix was transformed into a similarity matrix for network visualization and analysis (Friedman et al., 2012). The final adjacency matrices represented taxa as nodes and significant SparCC association coefficients as weighted edges, enabling both topological and functional analysis of microbial community structure across temperature gradients.

### 2.3. Network analysis

Constructed networks were analyzed using NetCoMi to calculate network summary metrics and identify key structural properties. Global network properties such as average path length, modularity, and clustering coefficient were calculated to characterize overall network structure (Peschel et al., 2021). To quantify network properties, betweenness centrality was used to identify hubs for each temperature. Hubs are nodes with a betweenness centrality above 95% of all nodes. Cohesion values (both positive and negative) were calculated for each sample (j) as the sum of significant positive or negative correlations between taxa, weighted by taxa abundances. The proportion of negative to positive cohesion values were calculated as the absolute value to examine high temperature stress (Hernandez et al., 2021). The networks were visualized in Gephi 0.10.1

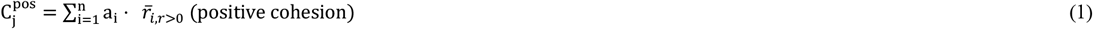

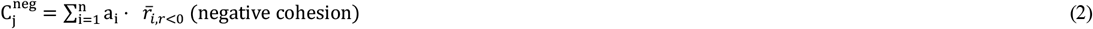

### 2.4. Statistical analysis

Each experimental sample included four independent biological replicates to ensure statistical robustness. For network construction, correlation analyses were performed with a significance threshold of p < 0.05, and only statistically significant correlations were included in the final networks. Microbial networks were visualized using Gephi software. Heatmaps were generated using the R package ComplexHeatmap. All figures were created and refined using Inkscape software.

## 3. Results and discussion

### 3.1. Temperature tolerant species and community restructuring under thermal stress

Metagenomic analysis revealed 140 species showing significant positive correlations with elevated temperatures (Fig. 1; r>0.5, p<0.05). Principal coordinate analysis (PCoA) based on Bray-Curtis distances demonstrated clear temperature-dependent community structuring, with samples from higher temperatures (32°C and 35°C) forming distinct clusters compared to moderate temperatures (23°C-29°C). This separation indicates substantial shifts in community composition under thermal stress. The emergence of 140 temperature-tolerant species represented a significant shift in community composition, suggesting the development of thermal adaptation strategies at the community level. This restructuring likely reflected both direct temperature effects and indirect effects through altered biogeochemical processes and resource availability. The clear separation of community structures at different temperatures, as demonstrated by PCoA analysis, indicated that temperature acted as a strong environmental filter, selecting species with appropriate thermal tolerance mechanisms. This selection pressure has led to the emergence of distinct community assemblages, with certain taxes showing remarkable adaptability across the temperature gradient.

**Fig. 1.**
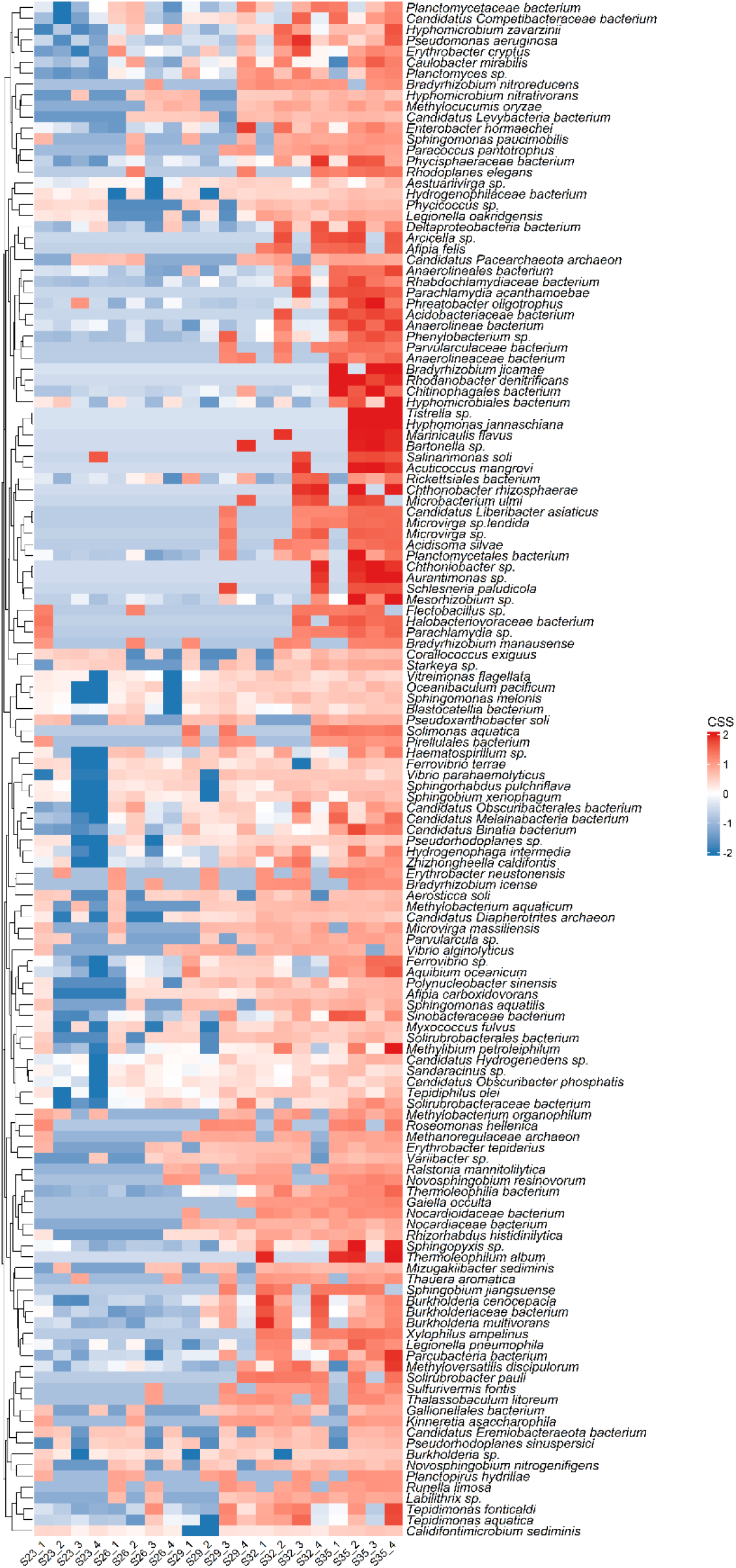
The abundance of 140 high temperature tolerant species.

### 3.2. Network Analysis of Microbial Interactions

Network analysis revealed significant changes in microbial community organization across temperature gradients. The network showed high modularity, with three distinct major modules emerging (Fig. 2): Module I: Dominated by thermotolerant species, particularly *Deltaproteobacteria*, which maintained presence across all temperature conditions; Module II: Characterized by moderate-temperature specialists; Module III: Comprised primarily of high-temperature specialists, including *Parcubacteria bacterium* and *Sphingobium xenophagum*. The species connected to tolerant species were extracted, and a network was constructed (Fig. 3). The entire network shows a general trend of decreasing nodes as temperature increases, from 1491 nodes at 23°C to 1181 nodes at 35°C, suggesting temperature stress reduces network complexity. However, the extracted network shows an opposite trend, with network size increasing from 57.48% at 23°C to 92.99% at 32°C of the entire network, indicating that higher temperatures lead to more significant interactions among the remaining species. In the entire network, the total number of edges decreases with an increasing temperature (121,904 at 23°C to 88,537 at 32°C), except for a slight increase at 35°C (94,633). The extracted network shows remarkably fewer edges (1.55-3.57% of the entire network), but the proportion increases with temperature, suggesting that while overall connections decrease, the remaining connections become more concentrated. In the entire network, the ratio of positive to negative edges remains relatively stable (1.04-1.13) across temperatures, maintaining a slight dominance of positive interactions. The extracted network shows a more pronounced positive bias, particularly at 23°C (1.61 positive/negative ratio), which gradually decreases to 1.00 at 32°C. This suggests that positive interactions are more crucial in the core network structure, especially at lower temperatures. The number of tolerant species increases substantially with temperature, from 34 at 23°C to 127 at 35°C, indicating adaptive responses to thermal stress. While the total number of hubs remains constant (116), the number of tolerant hubs increases from 8 at 23°C to 26 at 35°C, suggesting that key network positions are increasingly occupied by temperature-tolerant species. The maintenance of positive/negative edge ratios across temperatures in the entire network suggests an inherent stability mechanism. The increasing proportion of tolerant species and hubs indicates a selective pressure favoring temperature-resistant organisms, potentially representing an adaptive response to thermal stress.

**Fig. 2.**
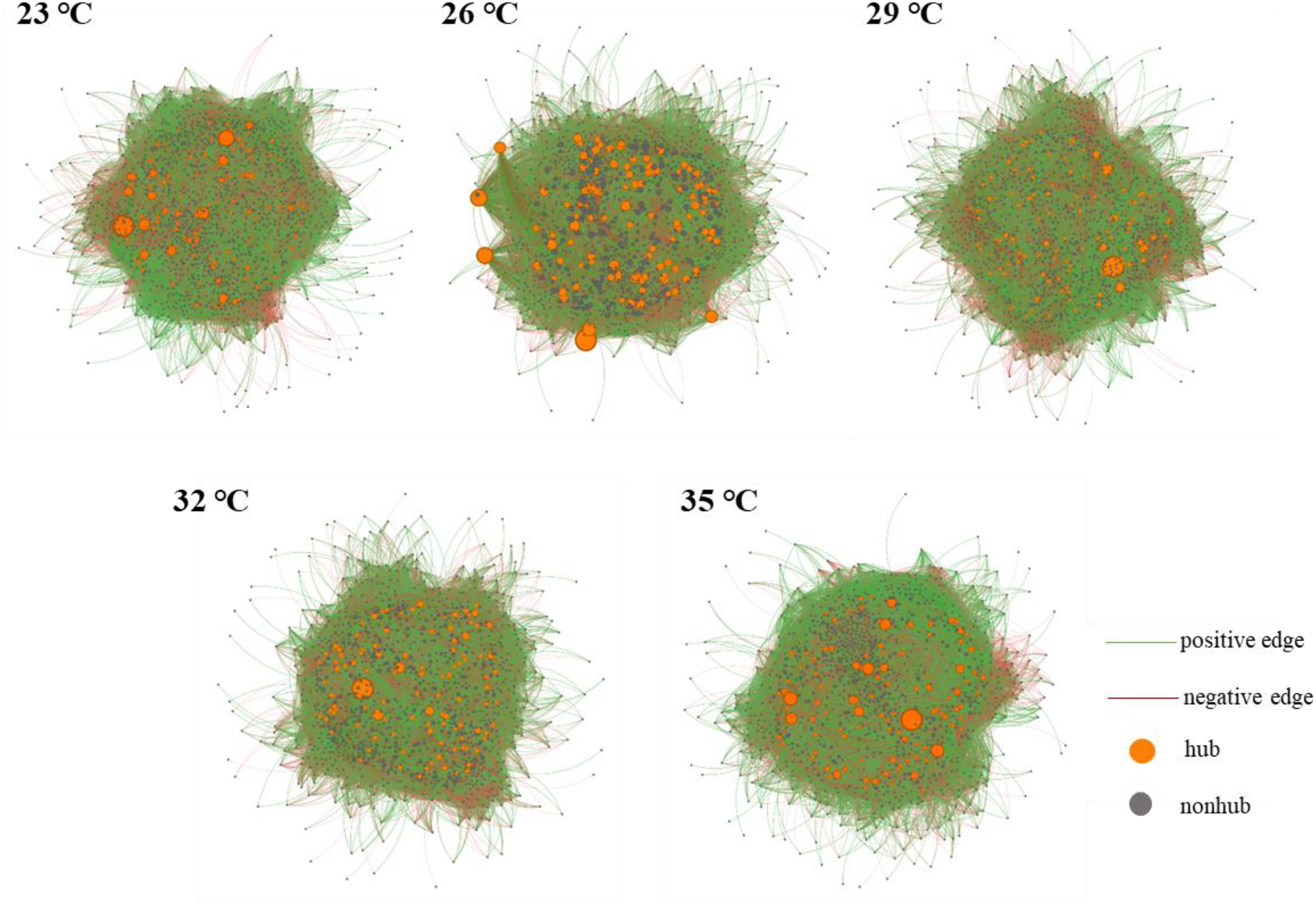
Microbial network across temperature gradients in Yellow River. Networks show all identified prokaryotic species at each temperature treatment (23°C, 26°C, 29°C, 32°C, and 35°C). Orange nodes represent hub species with high connectivity, and grey nodes represent non-hub species. Green edges indicate positive correlations between species pairs, while red edges indicate negative correlations.

**Fig. 3.**
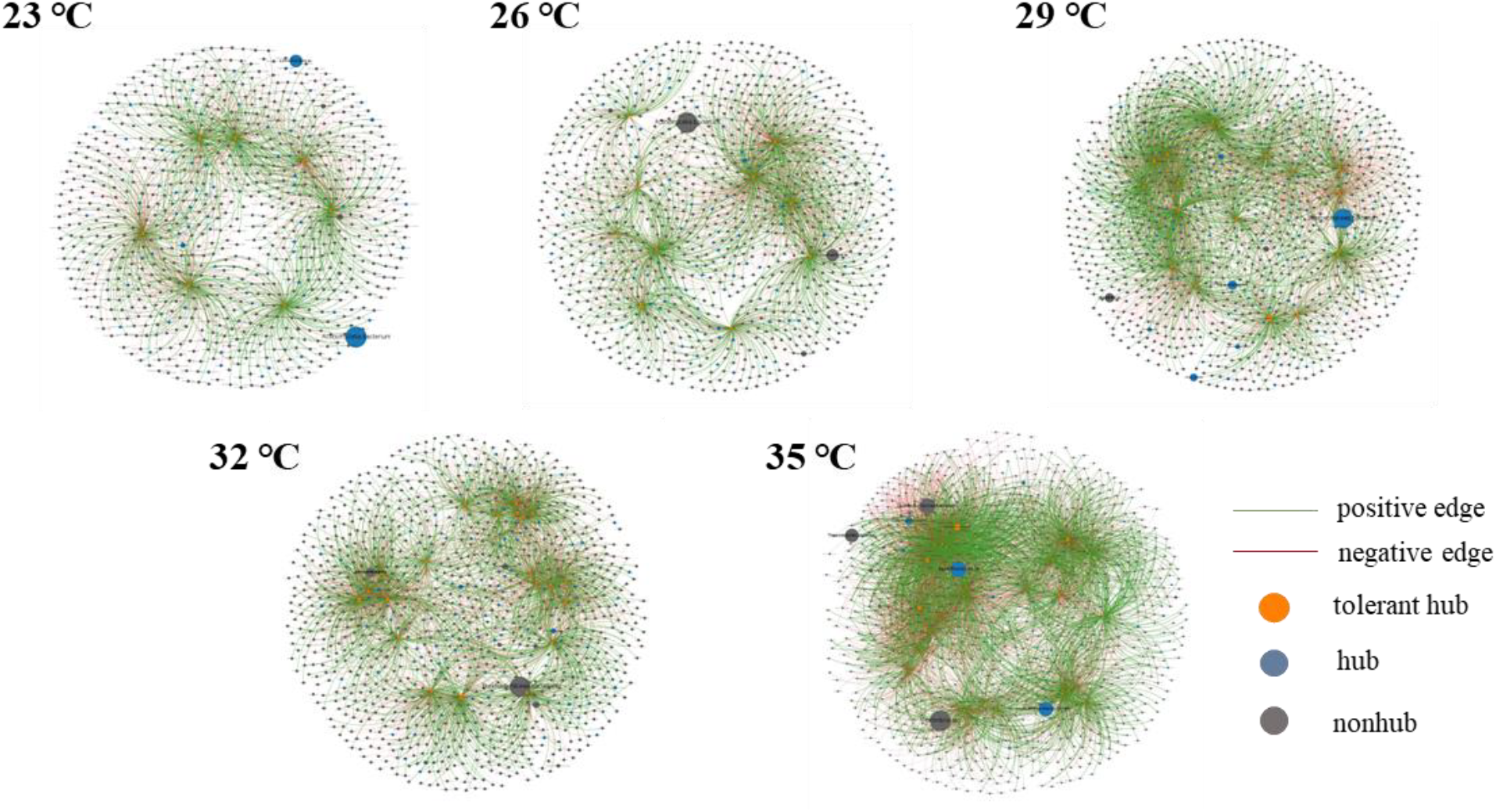
Extracted microbial networks across temperature gradients in Yellow River. Networks display all identified temperature tolerant hubs and their associated connections at each temperature treatment (23°C, 26°C, 29°C, 32°C, and 35°C). Orange nodes represent tolerant hub species that maintain high connectivity across temperature treatments, blue nodes represent hub species that are not temperature-tolerant, and grey nodes represent non-hub species. Green edges indicate positive correlations between species pairs, while red edges indicate negative correlations.

**Table 1.**
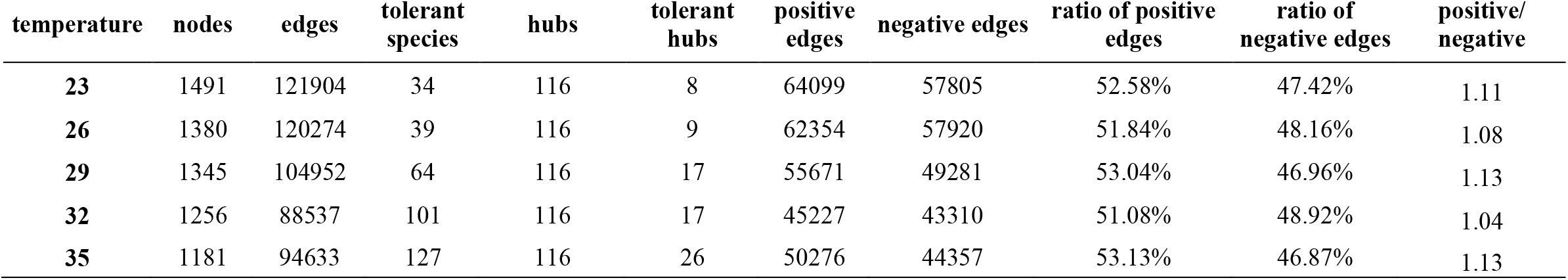
The properties of entire network.

| temperature | nodes | edges | tolerant species | hubs | tolerant hubs | positive edges | negative edges | ratio of positive edges | ratio of negative edges | positive/negative |
| --- | --- | --- | --- | --- | --- | --- | --- | --- | --- | --- |
| 23 | 1491 | 121904 | 34 | 116 | 8 | 64099 | 57805 | 52.58% | 47.42% | 1.11 |
| 26 | 1380 | 120274 | 39 | 116 | 9 | 62354 | 57920 | 51.84% | 48.16% | 1.08 |
| 29 | 1345 | 104952 | 64 | 116 | 17 | 55671 | 49281 | 53.04% | 46.96% | 1.13 |
| 32 | 1256 | 88537 | 101 | 116 | 17 | 45227 | 43310 | 51.08% | 48.92% | 1.04 |
| 35 | 1181 | 94633 | 127 | 116 | 26 | 50276 | 44357 | 53.13% | 46.87% | 1.13 |

### 3.3. Microbial Adaptations to Elevated Temperatures

These findings reveal significant adaptations in network structure and community dynamics in response to increasing temperatures (Pan et al., 2022). The network demonstrates a clear trend toward a more concentrated structure, evidenced by the increasing proportion of extracted network nodes relative to the entire network (from 57.48% at 23°C to 92.99% at 32°C), suggesting that while the overall network size decreases, the remaining interactions become more focused and potentially more crucial for community stability. This concentration is accompanied by a dramatic increase in temperature-tolerant species (from 34 to 127) and tolerant hubs (from 8 to 26), indicating a shift in community composition that favors thermally resistant organisms. Notably, despite this structural simplification, the network maintains a remarkable balance in its interaction types, with the positive-to-negative edge ratio remaining relatively stable (1.04-1.13) in the entire network, though showing more variation in the extracted network (1.61-1.00). This maintenance of interaction balance suggests an inherent stability mechanism that persists even as the network adapts to thermal stress. The community appears to develop thermal adaptation mechanisms, as evidenced by both the increasing proportion of tolerant species and the preservation of essential network properties. The retention of a constant number of hubs (116) while the proportion of tolerant hubs increases suggests a selective pressure that maintains network structure while favoring temperature-resistant species in key positions. This adaptation is further supported by the extracted network’s edge dynamics, where despite having fewer total edges (1.55-3.57% of the entire network), the proportion increases with temperature, indicating that while overall connections decrease, the remaining connections become more concentrated and potentially more significant for community function. These structural and compositional changes collectively suggest a sophisticated adaptive response to thermal stress, potentially providing insights into how similar ecological networks might respond to future climate change scenarios, where the maintenance of core network functions appears to be prioritized through the reorganization of species interactions and the emergence of thermally tolerant key players (Hernandez et al., 2021).

**Table 2.** The properties of extracted network.

| temperature | nodes | ratio of<br>extracted<br>to entire | edges | ratio of<br>extracted<br>to entire | tolerant hubs | positive<br>edges | ratio of positive<br>edges in total<br>edges | negative<br>edges | ratio of negative<br>edges<br>to total edges | positive/<br>negative |
| --- | --- | --- | --- | --- | --- | --- | --- | --- | --- | --- |
| 23 | 857 | 57.48% | 1888 | 1.55% | 8 | 1166 | 61.76% | 722 | 38.24% | 1.61 |
| 26 | 1126 | 81.59% | 2120 | 1.76% | 9 | 1134 | 53.49% | 986 | 46.51% | 1.15 |
| 29 | 1246 | 92.64% | 3617 | 3.45% | 17 | 1933 | 53.44% | 1684 | 46.56% | 1.15 |
| 32 | 1168 | 92.99% | 3162 | 3.57% | 17 | 1580 | 49.97% | 1582 | 50.03% | 1.00 |
| 35 | 1133 | 95.94% | 5256 | 5.55% | 26 | 2811 | 53.48% | 2445 | 46.52% | 1.15 |

## 4. Conclusions

This study reveals sophisticated microbial community adaptations to thermal stress through coordinated network restructuring and species composition shifts. Temperature acted as a strong environmental filter, selecting 140 thermotolerant species that increasingly dominated community structure as temperatures rose from 23°C to 35°C. Network analysis revealed a remarkable adaptive strategy characterized by a decrease in overall network complexity, with nodes dropping from 1491 to 1181. Despite this reduction, the remaining interactions became more concentrated and functionally important, as tolerant species increasingly occupied key hub positions, increasing from 8 to 26 tolerant hubs. The maintenance of interaction balance, with positive to negative edge ratios remaining stable at 1.04 to 1.13 despite structural simplification, suggests inherent stability mechanisms that persist under thermal stress. Three distinct modules emerged, with thermotolerant *Deltaproteobacteria* maintaining presence across all conditions while high-temperature specialists such as *Parcubacteria* and *Sphingobium xenophagum* dominated elevated temperature networks. These findings provide critical insights into microbial community resilience mechanisms under climate change scenarios. The selective pressure favoring temperature-resistant organisms in key network positions, combined with the preservation of essential interaction patterns, demonstrates how microbial communities can maintain functionality while adapting to thermal stress through strategic reorganization rather than complete restructuring. Understanding these adaptation mechanisms is crucial for predicting and managing the impacts of climate change on river ecosystems.

## Acknowledgements

The authors acknowledge the use of New Zealand eScience Infrastructure (NeSI) high performance computing facilities, as well as the consulting support and training services provided as part of this research. NeSI’s national facilities are jointly funded by NeSI’s collaborator institutions and the Ministry of Business, Innovation & Employment through the Research Infrastructure programme (https://www.nesi.org.nz). We also thank the Centre for eResearch at the University of Auckland for their valuable support in facilitating this work (http://www.eresearch.auckland.ac.nz). We are especially grateful to Dr. Qiaoling Yu and colleagues for making the metagenomics data publicly available under BioProject PRJ PRJEB54031 (http://www.ebi.ac.uk/ena/data/view/PRJEB54031), which has been immensely helpful in conducting this study.

## References

Aragaw, T. A., Bogale, F. M., & Gessesse, A. (2022). Adaptive Response of Thermophiles to Redox Stress and Their Role in the Process of dye Degradation From Textile Industry Wastewater. Frontiers in Physiology, 13. 10.3389/fphys.2022.908370

Baatout, S., Boever, P. D., & Mergeay, M. (2005). Temperature-induced changes in bacterial physiology as determined by flow cytometry. Ann. Microbiol.

Borisov, V. B., Siletsky, S. A., Nastasi, M. R., & Forte, E. (2021). ROS Defense Systems and Terminal Oxidases in Bacteria. Antioxidants, 10(6), Article 6. 10.3390/antiox10060839

Buchfink, B., Xie, C., & Huson, D. H. (2015). Fast and sensitive protein alignment using DIAMOND. Nature Methods, 12(1), 59–60. 10.1038/nmeth.3176

Hawkes, C. V., & Keitt, T. H. (2015). Resilience vs. historical contingency in microbial responses to environmental change. Ecology letters, 18(7), 612–625.

Hernandez, D. J., David, A. S., Menges, E. S., Searcy, C. A., & Afkhami, M. E. (2021). Environmental stress destabilizes microbial networks. The ISME Journal, 15(6), 1722–1734. 10.1038/s41396-020-00882-x

Hutchins, D. A., & Fu, F. (2017). Microorganisms and ocean global change. Nature Microbiology, 2(6), 17058.

Hutchins, D. A., Jansson, J. K., Remais, J. V., Rich, V. I., Singh, B. K., Trivedi, P. (2019). Climate change microbiology—problems and perspectives. Nature Reviews Microbiology, 17(6), 391–396.

Hyatt, D., Chen, G.-L., LoCascio, P. F., Land, M. L., Larimer, F. W., & Hauser, L. J. (2010). Prodigal: Prokaryotic gene recognition and translation initiation site identification. BMC Bioinformatics, 11(1), 119. 10.1186/1471-2105-11-119

Jones, A. (2022). The health impacts of climate change: Why climate action is essential to protect health. Orthopaedics and Trauma, 36(5), 248–255. 10.1016/j.mporth.2022.07.001

Li, D., Luo, R., Liu, C.-M., Leung, C.-M., Ting, H.-F., Sadakane, K., Yamashita, H., & Lam, T.-W. (2016). MEGAHIT v1.0: A fast and scalable metagenome assembler driven by advanced methodologies and community practices. Methods, 102, 3–11. 10.1016/j.ymeth.2016.02.020

Linzner KA, Kent AG and Martiny AC (2018) Evolutionary Pathway Determines the Stoichiometric Response of Escherichia coli Adapted to High Temperature. Front. Ecol. Evol. 5:173.

Mise, K., & Iwasaki, W. (2022). Unexpected absence of ribosomal protein genes from metagenome-assembled genomes. ISME Communications, 2(1), 118. 10.1038/s43705-022-00204-6

Moldogazieva, N. T., Mokhosoev, I. M., Feldman, N. B., & Lutsenko, S. V. (2018). ROS and RNS signalling: Adaptive redox switches through oxidative/nitrosative protein modifications. Free Radical Research, 52(5), 507–543. 10.1080/10715762.2018.1457217

Nakagawa, S., Takai, K., Inagaki, F., Hirayama, H., Nunoura, T., Horikoshi, K., & Sako, Y. (2005). tDistribution, phylogenetic diversity and physiological characteristics of epsilon-proteobacteria in a deep-sea hydrothermal field. Environmental Microbiology, 7(11), 1619–1632.

Obeagu, E. (2018). A Review on Free Radicals and Antioxidants. 4, 123–133. 10.22192/ijcrms.2018.04.02.019

Pan, S., Zhu, C., Zhao, X.-M., & Coelho, L. P. (2022). A deep siamese neural network improves metagenome-assembled genomes in microbiome datasets across different environments. Nature Communications, 13(1), 2326. 10.1038/s41467-022-29843-y

Paulson, J. N., Stine, O. C., Bravo, H. C., & Pop, M. (2013). Differential abundance analysis for microbial marker-gene surveys. Nature Methods, 10(12), 1200–1202. 10.1038/nmeth.2658

Pecl, G. T., Araújo, M. B., Bell, J. D., Blanchard, J., Bonebrake, T. C., Chen, I. C., … & Williams, S. E. (2017). Biodiversity redistribution under climate change: Impacts on ecosystems and human well-being. Science, 355(6332), eaai9214.

Peschel, S., Müller, C. L., Von Mutius, E., Boulesteix, A. L., & Depner, M. (2021). NetCoMi: network construction and comparison for microbiome data in R. Briefings in bioinformatics, 22(4), bbaa290.

Peschel, S., Müller, C. L., von Mutius, E., Boulesteix, A.-L., & Depner, M. (2021). NetCoMi: Network construction and comparison for microbiome data in R. Briefings in Bioinformatics, 22(4), bbaa290. 10.1093/bib/bbaa290

Poloczanska, E.S., Brown, C.J., Sydeman, W.J., Kiessling, W., Schoeman, D.S., Moore, P.J., … Duarte, C.M. (2013). Global imprint of climate change on marine life. Nature Climate Change, 3(10), 919–925.

Ren, L., Lu, Z., Xia, X., Peng, Y., Gong, S., Song, X., … & Wu, Q. L. (2022). Metagenomics reveals bacterioplankton community adaptation to long-term thermal pollution through the strategy of functional regulation in a subtropical bay. Water Research, 216, 118298.

Reusch, T. B. (2014). Climate change in the oceans: evolutionary versus phenotypically plastic responses of marine animals and plants. Evolutionary Applications, 7(1), 104–122.

Savo, V., Lepofsky, D., Benner, J. P., Kohfeld, K. E., Bailey, J., & Lertzman, K. (2016). Observations of climate change among subsistence-oriented communities around the world. Nature Climate Change, 6(5), 462–473.

Sewe, S. O., Silva, G., Sicat, P., Seal, S. E., & Visendi, P. (2022). Trimming and Validation of Illumina Short Reads Using Trimmomatic, Trinity Assembly, and Assessment of RNA-Seq Data. In D. Edwards (Ed.), Plant Bioinformatics: Methods and Protocols (pp. 211–232). Springer US. 10.1007/978-1-0716-2067-0_11

Singh, N., Singh, J., & Singh, K. (2018). Small at Size, Big at Impact: Microorganisms for Sustainable Development. In J. Singh, D. Sharma, G. Kumar, & N. R. Sharma (Eds.), Microbial Bioprospecting for Sustainable Development (pp. 3–28). Springer. 10.1007/978-981-13-0053-0_1

Song, N., Wu, D., Xu, H., & Jiang, H. (2022). Integrated evaluation of the reactive oxygen species (ROS) production characteristics in one large lake under alternating flood and drought conditions. Water Research, 225, 119136.

Suzuki, N., & Mittler, R. (2006). Reactive oxygen species and temperature stresses: A delicate balance between signaling and destruction. Physiologia Plantarum, 126(1), 45–51. 10.1111/j.0031-9317.2005.00582.x

Tamames, J., & Puente-Sánchez, F. (2019). SqueezeMeta, a highly portable, fully automatic metagenomic analysis pipeline. Frontiers in microbiology, 9, 3349.

Tamames, J., & Puente-Sánchez, F. (2019). SqueezeMeta, A Highly Portable, Fully Automatic Metagenomic Analysis Pipeline. Frontiers in Microbiology, 9. 10.3389/fmicb.2018.03349

Tsilimigras, M. C. B., & Fodor, A. A. (2016). Compositional data analysis of the microbiome: Fundamentals, tools, and challenges. Annals of Epidemiology, 26(5), 330–335. 10.1016/j.annepidem.2016.03.002

Tsilimigras, M. C., & Fodor, A. (2016). Compositional data analysis of the microbiome: fundamentals, tools, and challenges. Annals of epidemiology, 26(5), 330–335.

Uritskiy, G., & DiRuggiero, J. (2019). Applying Genome-Resolved Metagenomics to Deconvolute the Halophilic Microbiome. Genes, 10(3), Article 3. 10.3390/genes10030220

Xiao, W., & Loscalzo, J. (2020). Metabolic Responses to Reductive Stress. Antioxidants & Redox Signaling, 32(18), 1330–1347. 10.1089/ars.2019.7803

Yeh, Y. C., & Fuhrman, J. A. (2022). Effects of phytoplankton, viral communities, and warming on free-living and particle-associated marine prokaryotic community structure. Nature Communications, 13(1), 7905.

Yu, M., Eglinton, T. I., Haghipour, N., Montluçon, D. B., Wacker, L., Hou, P., Zhang, H., & Zhao, M. (2019). Impacts of Natural and Human-Induced Hydrological Variability on Particulate Organic Carbon Dynamics in the Yellow River. Environmental Science & Technology, 53(3), 1119–1129. 10.1021/acs.est.8b04705

Yu, Q., Feng, T., Yang, J., Su, W., Zhou, R., Wang, Y., Zhang, H., & Li, H. (2022). Seasonal distribution of antibiotic resistance genes in the Yellow River water and tap water, and their potential transmission from water to human. Environmental Pollution, 292, 118304. 10.1016/j.envpol.2021.118304

Yu, Q., Han, Q., Shi, S., Sun, X., Wang, X., Wang, S., Yang, J., Su, W., Nan, Z., & Li, H. (2023). Metagenomics reveals the response of antibiotic resistance genes to elevated temperature in the Yellow River. Science of The Total Environment, 859, 160324. 10.1016/j.scitotenv.2022.160324

Yu, Y., Liu, L., Wang, J., Zhang, Y., & Xiao, C. (2021). Effects of warming on the bacterial community and its function in a temperate steppe. Science of The Total Environment, 792, 148409. 10.1016/j.scitotenv.2021.148409

Zhang, M., Wu, J., Sun, R., Tao, X., Wang, X., Kang, Q., Wang, H., Zhang, L., Liu, P., Zhang, J., Xia, Y., Zhao, Y., Yang, Y., Xiong, Y., Guan, K.-L., Zou, Y., & Ye, D. (2019). SIRT5 deficiency suppresses mitochondrial ATP production and promotes AMPK activation in response to energy stress. PLOS ONE, 14(2), e0211796. 10.1371/journal.pone.0211796

Zhao, M. M., Chen, Y., Xue, L., Fan, T. T., & Emaneghemi, B. (2018). Greater health risk in wet season than in dry season in the Yellow River of the Lanzhou region. Science of The Total Environment, 644, 873–883. 10.1016/j.scitotenv.2018.07.006

Zheng, J., Cao, J., Mao, Y., Su, Y., & Wang, J. (2019). Comparative transcriptome analysis provides comprehensive insights into the heat stress response of Marsupenaeus japonicus. Aquaculture, 502, 338–346. 10.1016/j.aquaculture.2018.11.023

Zhou, Y., Sun, B., Xie, B., Feng, K., Zhang, Z., Zhang, Z., Li, S., Du, X., Zhang, Q., Gu, S., Song, W., Wang, L., Xia, J., Han, G., & Deng, Y. (2021). Warming reshaped the microbial hierarchical interactions. Global Change Biology, 27(24), 6331–6347. 10.1111/gcb.15891

